# Primary cilia promote cardiac fibrosis and limit heart function after myocardial infarction

**DOI:** 10.64898/2026.05.30.728594

**Authors:** Xinyue Liu, Alessandra Norris, Ambili Appu, Emily Wilson, Hao Zhang, Jeffrey Olgin, Jeremy Reiter, Daniel Kopinke

## Abstract

Cardiomyocytes die and do not regenerate after an injury such as a myocardial infarction (MI), a leading cause of mortality worldwide. Following MI, cardiac fibroblasts (CFs) proliferate and differentiate into myofibroblasts, which then produce increased collagen and extracellular matrix (ECM) leading to fibrosis. Fibrosis can weaken cardiac output via excessive stiffening and interference with electric signal transmission, but can also prevent wall rupture under load (reviewed in (1)). Thus, dampening fibrosis has been investigated as a potential therapeutic intervention. Most mammalian cells possess a single primary cilium involved in intercellular communication. We investigated the role of CF primary cilia in sensing injury signals and initiating fibrotic remodeling. We found that deleting CF cilia reduced fibrosis and improved cardiac output after MI, demonstrating that cilia act as a signaling hub that amplifies the fibrotic response in the injured heart.

---

Within the heart, CFs are the main ciliated cell type and accumulate in injured areas after myocardial injury (2). To test the *in vivo* function of CF primary cilia during cardiac injury responses, we created an inducible conditional mouse model that removes cilia from CFs upon tamoxifen administration. More specifically, we generated *Pdgfra*^*CreERT*^ *Ift88*^*lox/lox*^ mice that, upon tamoxifen administration, delete *Ift88*, a gene required for ciliogenesis and ciliary maintenance, in CFs (Figure 1A). Consequently, we refer to tamoxifen-treated *Pdgfra*^*CreERT*^ *Ift88*^*lox/lox*^ mice as CF^no cilia^ mice, hereafter.

**Figure 1.**
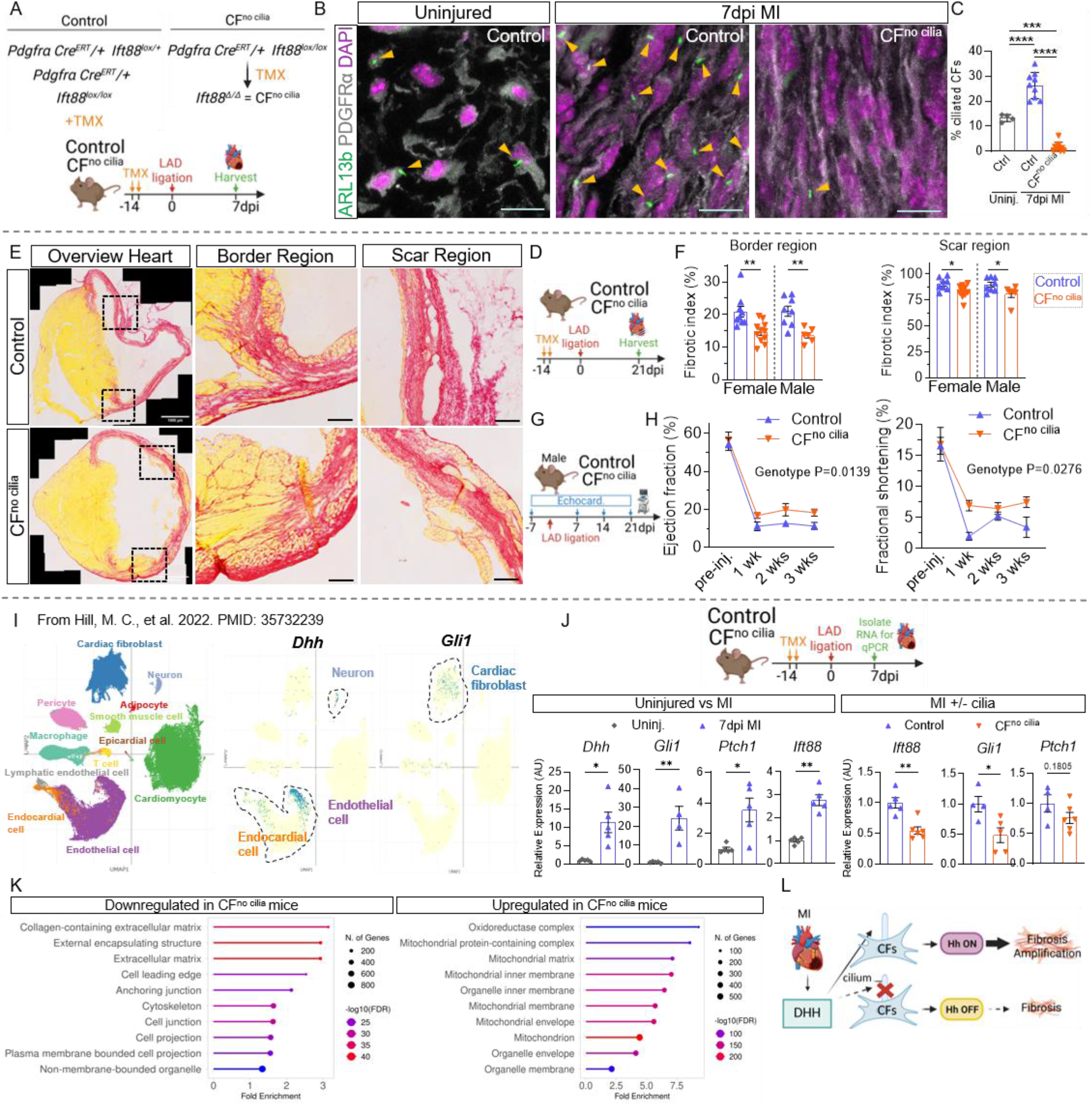
Loss of primary cilia within CFs results in reduced fibrosis and improved heart function. **(A)** Genetic scheme and experimental outline for B, C. **(B)** Representative images of 1-year-old uninjured wild-type mice, 7 days post-LAD surgery control and CF^no cilia^ mice, with immunofluorescent staining for ARL13B (cilia, arrows), PDGFRα (CFs) and DAPI (nuclei). Scale bar: 10 µm. **(C)** Percentage of ciliated CFs. n = 5–8 mice per group. Data represent mean ± SEM. Unpaired two-tailed Student’s t-test. **(D)** Experimental outline for D, F. **(E)** Sirius red staining of control and CF^no cilia^ hearts, 21 days post myocardial infarction. Left: cross-section of the whole heart, scale bar: 1000 µm. Middle: magnified view of the infarct border, scale bar: 10 µm. Left: magnified view of the post-infarct scar, scale bar: 10 µm. n = 8–12 mice per group. **(F)** Collagen content (%) in the infarct border and scar of control and CF^no cilia^ hearts, 21 days post-myocardial infarction. n = 8–12 mice per group per sex. Data represent means ± SEM. Unpaired two-tailed Student’s t-test. **(G)** Experimental outline for H. **(H)** Ejection fraction and fractional shortening of control and CF^no cilia^ hearts a week prior to injury (pre-injury), and 1-, 2- and 3-weeks post injury. n = 8–10 male mice per group. Data represent mean ± SEM. Multiple t-tests. **(I)** UMAP (Uniform Manifold Approximation and Projection) plot of published single-nucleus RNA sequencing on 157,273 nuclei from hearts of control and congenital heart disease patients. **(J)** RT-qPCR for *Dhh, Gli1, Ptch1* and *Ift88* from wild-type heart ventricles of 10-week-old uninjured mice and 7 days post-injury. RT-qPCR for *Ift88, Gli1* and *Ptch1* from control and CF^no cilia^ heart ventricles 7 days post-injury. n = 4–6 mice per group. Data represent mean ± SEM. Unpaired two-tailed Student’s t-test. Expression normalized to housekeeping genes *Hprt, Sra1* and *Pde12*. **(K)** Bulk RNA sequencing of control and CF^no cilia^ hearts 7 dpi. CF^no cilia^ mice show reduced expression of genes associated with fibrosis (*e*.*g*., *Col1a1/2, Col3a1, Postn, Ctgf, Fn1, Lox*) and inflammation *(e*.*g*., *Il1b, S100a8/9, Lgals3*), and increased expression of genes associated with mitochondrial energy pathways. **(L)** Schematic depicting how, after MI, DHH signals through CF primary cilia to activate Hh-dependent fibrosis.

We induced MI by ligating the descending artery in control and CF^no cilia^ mice and analyzed heart histology 7 days post injury (dpi) (Figure 1A). Cardiac injury increased ciliation in CFs at 7dpi in control animals (Figure 1B and C), consistent with a role for CF cilia in cardiac remodeling. CF ciliation in CF^no cilia^ mice was abrogated, as expected (Figure 1B and C).

To test whether loss of CF cilia impacts cardiac fibrosis post-MI, we blindly evaluated collagen deposition through a Sirius Red staining in control and CF^no cilia^ mice 21dpi MI (Figure 1D and E). In both sexes, collagen in CF^no cilia^ hearts was decreased compared to controls (Figure 1F). Collagen deposition was decreased at both the border and scar regions of CF^no cilia^ hearts. Thus, removal of primary cilia from CFs decreases fibrosis after MI, indicating that cilia amplify the fibrotic response upon injury. Consistent with our *in vivo* findings, cilia promote fibrogenesis *in vitro* (2).

We also assessed whether CF cilia affect recovery of cardiac function post MI. We measured ejection fraction and fractional shortening by echocardiogram pre-injury and at 7, 14 and 21 days post MI in control and CF^no cilia^ male mice (Figure 1G). Prior to injury, control and CF^no cilia^ hearts exhibited equivalent ejection fraction and fractional shortening (Figure 1H). Following MI, CF^no cilia^ hearts exhibited higher rejection fraction and fractional shortening compared to control hearts (Figure 1H). Thus, CF^no cilia^ hearts develop both decreased fibrosis and increased cardiac function post-MI.

To test whether the improved cardiac output was due to changes in cardiomyocyte density and/or size, we histologically evaluated post-MI cardiomyocytes. Control and CF^no cilia^ cardiomyocytes were of equal density and size 21 dpi (Figure S1A). Additionally, cardiomyocyte size distributions were similar across three cardiac regions (Figure S1B). These findings suggest that the increased cardiac function observed in CF^no cilia^ hearts post-MI is due to dampening of fibrosis.

Among the intercellular signals transduced by primary cilia is Hedgehog (Hh). Previously, we showed that ciliary Hh signaling regulates fibroblast cell fate after skeletal muscle injury, specifically through the Hh ligand, Desert Hedgehog (DHH) (3,4). To determine whether Hh signaling might amplify fibrosis in the heart, we queried publicly available single-nucleus RNA sequencing datasets from healthy human hearts and patients with congenital heart disease (5) (Figure 1I), as well as a murine single-cell RNA sequencing database of a pro-fibrotic environment, induced by administration of angiotensin II (6) (Figure S2A). These data indicated that Schwann and endothelial cells robustly express *DHH* (Figure 1I and S2A). In contrast, the genes encoding the other two Hh ligands (Sonic Hedgehog and Indian Hedgehog) were minimally expressed (Figure S2B and C). These data further indicated that the cells most robustly expressing the Hh target gene *GLI1* were cardiac fibroblasts (Figure 1I and S2A). These data are consistent with Schwann and endothelial cells robustly activating Hh signaling in CFs via DHH.

To determine whether Hh signaling is activated post MI, we evaluated gene expression of *Dhh, Ift88, Gli1* and *Ptch1*, another Hh target gene, via RT-qPCR 7dpi post MI. The expression of all these genes was increased post injury (Figure 1J). Thus, MI strongly induces Hh pathway activation.

To determine if loss of cilia blunts Hh pathway activation, we assessed Hh pathway activity in CF^no cilia^ mice 7dpi. Expression of *Ift88, Gli1* and *Ptch1* decreased in CF^no cilia^ mice compared to controls (Figure 1J). Taken together, these data demonstrate that cardiac injury induces Hh signaling via DHH, which is sensed by CF cilia to induce fibrosis.

We evaluated the broader transcriptional consequences of primary cilia loss in CFs by carrying out bulk RNA sequencing on control and CF^no cilia^ hearts 7dpi (Figure 1K). 420 genes were differentially expressed in CF^no cilia^ post-MI hearts as compared to control post-MI hearts (Figure S2D). Gene ontology (GO) enrichment analysis indicated that extracellular matrix and structural remodeling genes were downregulated in CF^no cilia^ post-MI hearts (Figure 1K). Mitochondrial and metabolic genes were upregulated in CF^no cilia^ post-MI hearts (Figure 1K). Together, these findings support the histological finding that removing primary cilia from CFs attenuates fibrosis following MI.

To assess whether removing CF cilia had long-term effects on heart homeostasis, we analyzed CF^no cilia^ and control hearts 14 months after tamoxifen administration (Figure S2A). CF^no cilia^ and control hearts exhibited no differences in collagen deposition (Figure S2B) or cardiomyocyte size (Figure S2C). Thus, CF cilia are not critical for cardiac homeostasis. Instead, they amplify the CF fibrotic response after MI and contribute to reduced cardiac function. Targeting cilia and/or Hh signaling to modulate the fibrotic response could provide a valuable strategy to improve cardiac outcomes following MI.

## SUPPLEMENTARY INFORMATION

**Supplementary Figure 1.**
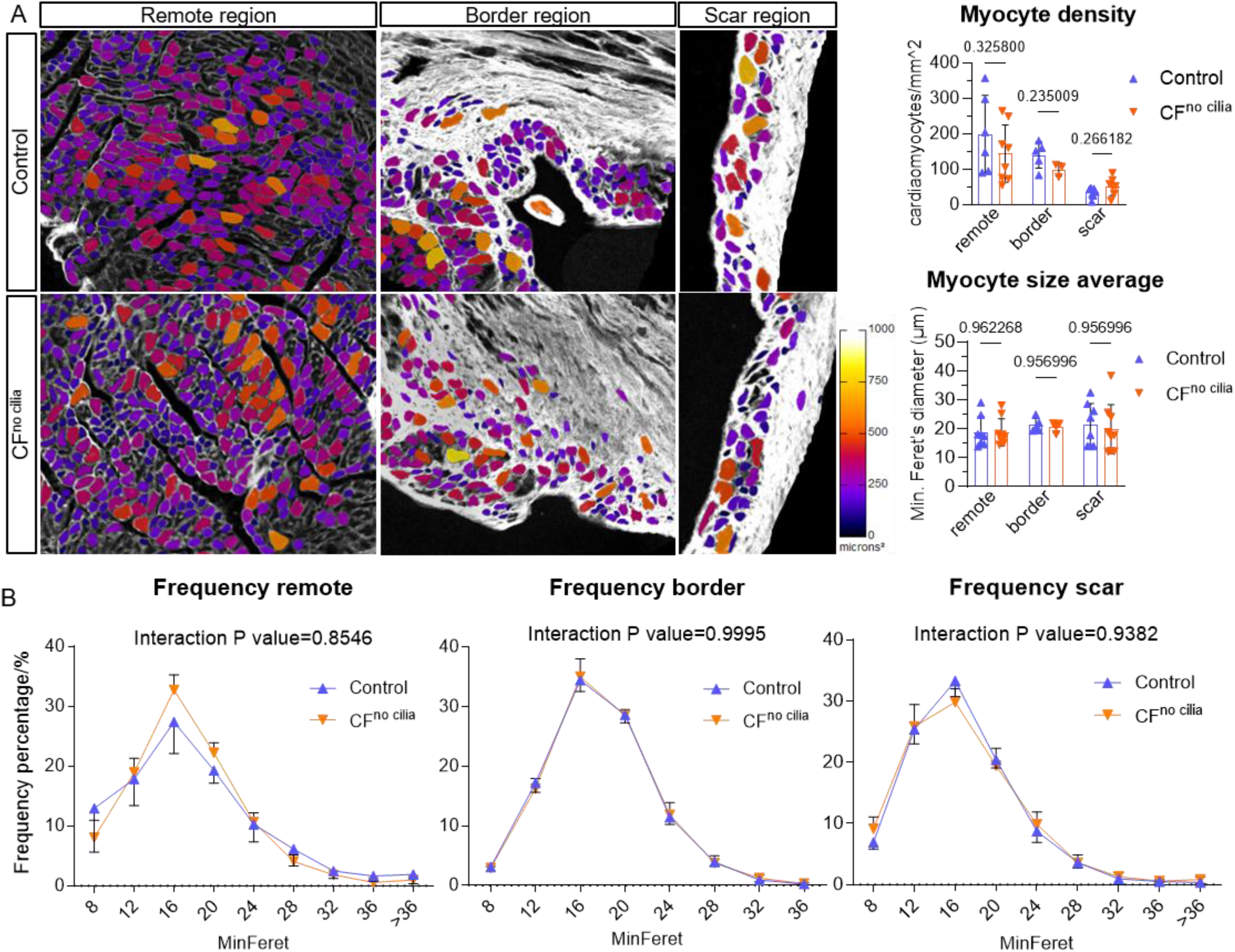
Cardiac fibroblast-specific cilia knockout does not impact cardiac myofiber size 21 days after MI. **(A)** Representative cross-sectional images, cardiomyocyte size and density quantifications of areas remote to, on the border of, and within the scar of control and CF^no cilia^ hearts after MI. Images are color-coded by individual cardiomyocyte size, as indicated in the scale (µm). n = 6 per group. **(B)** Cardiomyocyte size frequency distributions of the remote, border, and scar regions from control and CF^no cilia^ heart cross-sections. n = 6 per group. Data are presented as mean ± SEM. (A) Unpaired two-tailed Student’s t-test. (B) Two-way ANOVA with Tukey’s multiple comparisons test.

**Supplementary Figure 2.**
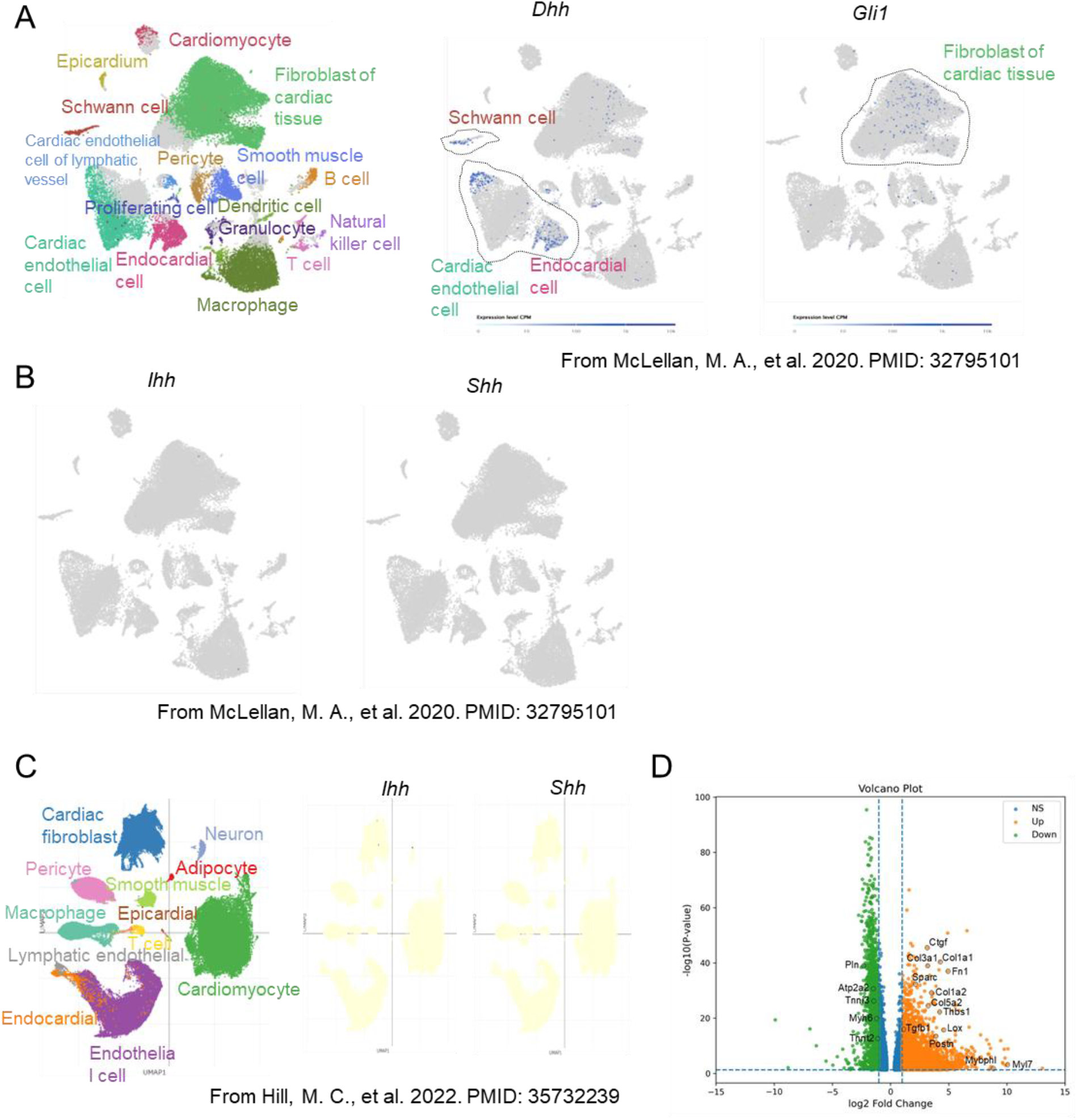
*Dhh* is the predominant Hh gene expressed in the heart and CF cilia loss attenuates the pro-fibrotic transcriptional program after MI. **(A)** UMAP (Uniform Manifold Approximation and Projection) plots from a published single-cell RNA sequencing dataset of chronic cardiac stress showing cell type annotations and log-normalized expression of *Dhh* and *Gli1* across cell clusters. *Dhh* is expressed in Schwann and endothelial cells. *Gli1* is expressed in cardiac fibroblasts. Data reanalyzed from McLellan, M. A., et al. 2020 (PMID: 32795101). **(B)** UMAP plots from the same chronic stress dataset (PMID: 32795101) showing log-normalized expression of *Ihh* and *Shh*. Neither is significantly expressed in the heart under chronic stress conditions. **(C)** UMAP plots from a published single-cell RNA sequencing dataset of congenital heart disease showing cell type annotations and log-normalized expression of *Ihh* and *Shh*. Neither is significantly expressed. Data reanalyzed from Hill, M. C., et al. 2022 (PMID: 35732239). **(D)** Volcano plot of differentially expressed genes from bulk RNA sequencing of CF^no cilia^ and control hearts at 7 days post-myocardial infarction. Upregulated (Up) and downregulated (Down) genes are in orange and green, respectively; select pro-fibrotic and inflammatory genes are labeled.

**Supplementary Figure 3.**
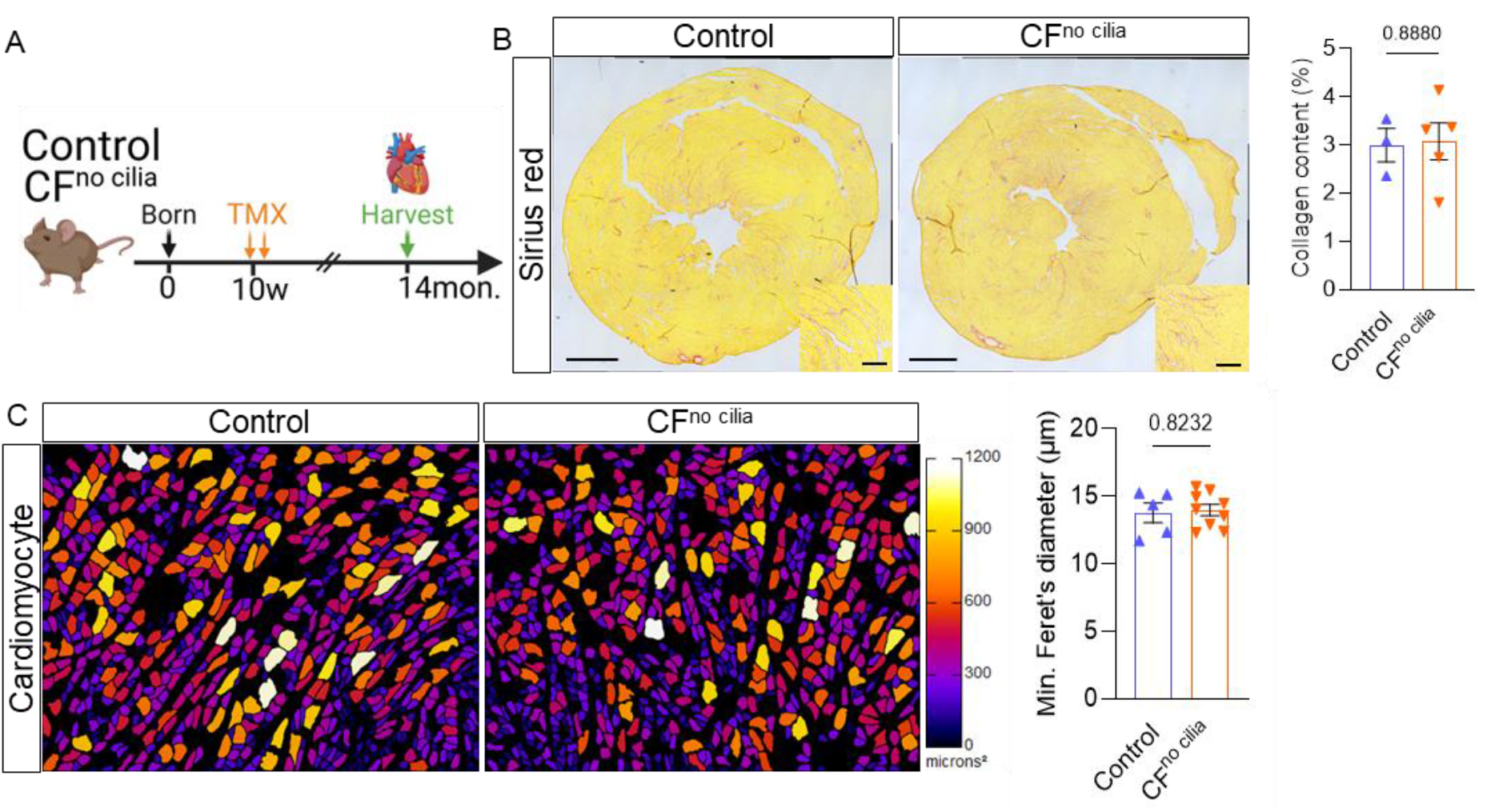
Removal of CF cilia does not significantly impact fibrosis or cardiomyocyte size under homeostatic conditions. **(A)** Experimental outline. **(B)** Representative cross-sectional images (scale bar: 500 µm; magnified images scale bar: 100 µm). Sirius red staining quantification of 14-month-old control and CF^no cilia^ hearts. **(C)** Representative cross-sectional images and cardiomyocyte size quantification of control and CF^no cilia^ mice after MI. Images are color-coded by individual cardiomyocyte size, as indicated in the scale (µm). n = 5–6 per group (B); n = 5–8 per group (C). Data are presented as mean ± SEM. Unpaired two-tailed Student’s t-test.

**Supplementary Table 1.**
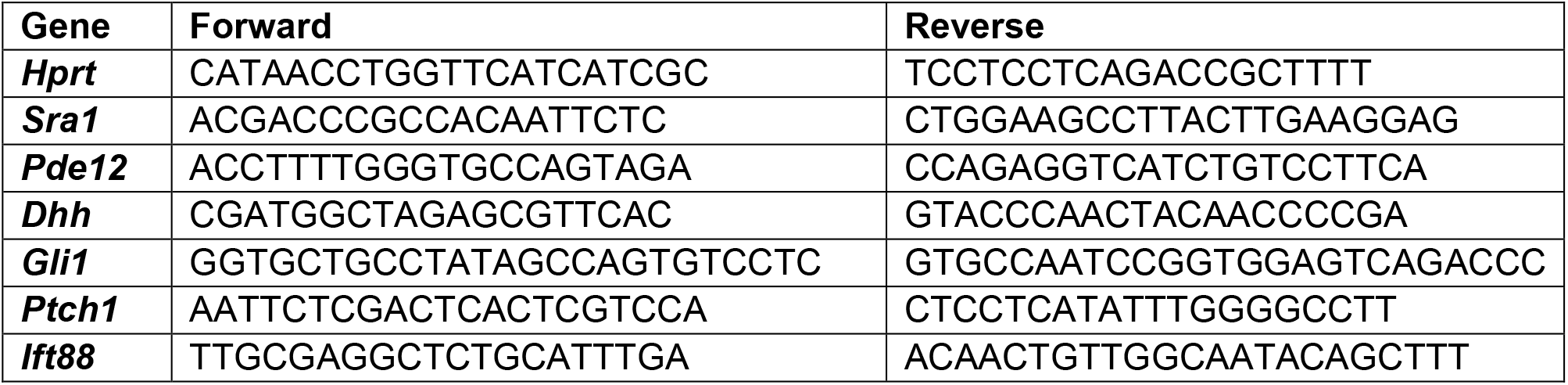
The qPCR primers used in the experiments.

## Materials and Methods

### Sex as a biological variable

To account for sex as a biological variable, male and female subjects were analyzed separately for all key phenotypes. Echocardiographic assessments were performed in male subjects only. Data for each sex are presented independently to ensure transparency and to facilitate sex-disaggregated interpretation of the results.

### Animal Studies

All mouse alleles used in this study have been previously reported. For conditional deletion of IFT88 specifically from CFs, a tamoxifen-inducible *Pdgfra*^*CreERT2*^ allele (Jax# 032770) was crossed to the *Ift88*^tm1Bky^ allele (Jax# 022409) and maintained on the CD1 background. PDGFRα is the gold-standard marker for identifying CFs in both murine and human muscle. To induce deletion of target genes, Tamoxifen (TRC, T006000) was dissolved in corn oil and administered via oral gavage at 200–250 mg/kg on two consecutive days. A two-week recovery period was observed following tamoxifen administration prior to injury induction. All mice, regardless of genotype, were subjected to tamoxifen gavage. Littermates lacking the Cre allele or heterozygous for the floxed allele were used as controls. Mice were housed in standard ventilated cages under controlled conditions (22–23°C, 40–50% humidity, 12-hour light/dark cycle) with ad libitum access to food and water. All animal procedures were approved by the Institutional Animal Care and Use Committee (IACUC) of the University of Florida and the University of California, San Francisco.

### Left anterior descending (LAD) coronary artery ligation injury model

MI was induced by permanent ligation of the LAD coronary artery. Briefly, mice were anesthetized with 1.0–2.0% isoflurane, intubated, and ventilated using a rodent ventilator. The chest was opened by left thoracotomy at the fourth intercostal space, and the pericardium was gently removed to expose the heart. The LAD coronary artery was identified and permanently ligated using a 6-0 silicone coated braided silk suture approximately 1–2 mm below the left atrial appendage. Successful ligation was confirmed by immediate visual blanching and cyanosis of the anterior left ventricular wall distal to the ligation site. The chest was then closed in layers, and mice were allowed to recover on a heated pad. Age-matched uninjured mice were used as controls. Surgeons were blinded to the genotypes.

### Echocardiograph

Cardiac function was assessed by transthoracic B-mode echocardiography at pre-surgery baseline (day −7), and at 7, 14, and 21 days following left anterior descending (LAD) coronary artery ligation. Mice were lightly anesthetized with 1.0–2.0% isoflurane and placed in a supine position on a heated platform to maintain body temperature. Images were acquired in the parasternal short-axis view at the level of the papillary muscles using the Vevo 3100 system (FUJIFILM VisualSonics, Toronto, Canada) equipped with a high-frequency transducer. Left ventricular internal diameter at end-diastole (LVID;d) and end-systole (LVID;s) were measured by tracing the endocardial border on 2D B-mode images. Ejection fraction (EF) and fractional shortening (FS) were calculated using the Vevo LAB software. A minimum of three consecutive cardiac cycles was averaged per measurement. All acquisitions and analyses were performed by a single operator blinded to experimental group allocation.

### Histology, Immunohistochemistry and Image Analysis

Briefly, upon harvesting, heart ventricles were bisected: the proximal third was used for RNA isolation, while the distal two-thirds were used for histology and immunostaining. Tissues were fixed in 4% paraformaldehyde (PFA) for 2.5 hours at 4°C, washed, and cryoprotected overnight in 30% sucrose. Ventricles were placed in OCT-filled cryomolds (Sakura, cat. no. 4566) and snap-frozen in isopentane cooled with liquid nitrogen. Three to four cryosections per ventricle were collected using a Leica cryostat at 10–12 µm thickness at intervals of 250–350 µm. Sections were incubated with primary antibodies in blocking solution (5% donkey serum in PBS containing 0.3% Triton X-100) overnight at 4°C. Primary antibodies used were as follows: rabbit anti-ARL13B (1:1000; Proteintech, cat. no. 17711-1-AP), rabbit anti-Laminin (1:1000; Sigma-Aldrich, cat. no. L9393), and goat anti-PDGFRα (1:250; R&D Systems, cat. no. AF1062). Alexa Fluor-conjugated secondary antibodies (Life Technologies, 1:1000) were applied in combination with Phalloidin-Alexa Fluor 568 (1:200; Molecular Probes, cat. no. A12380) for 1 hour at room temperature, after which slides were mounted using fluorescence mounting medium (SouthernBiotech, cat. no. 0100-01). Nuclei were counterstained with DAPI (Invitrogen, cat. no. D1306). Fibrillar collagen was visualized using Picrosirius Red staining, and Sirius Red-positive areas were quantified using the Color Threshold function in ImageJ. Images from randomly selected fields across multiple sections (each 250–350 µm apart) were acquired using a Leica DMi8 microscope equipped with an SPE confocal module and a high-resolution color camera. Whole ventricle cross-sectional images were generated using the Navigator function within the Leica LAS software. All images were processed identically and quantified using Fiji/ImageJ (v1.52p).

To evaluate the cross-sectional area (CSA) of injured and uninjured hearts, 5–7 images at 10× magnification per mouse were acquired in regions where myofibers were sectioned perpendicularly. Sections were stained with PHALLOIDIN or LAMININ to visualize myofibers. Uninjured areas, defined as regions in which myofibers lacked centrally located nuclei, were excluded from analysis during image processing in ImageJ. Individual myofibers were segmented using Cellpose, an AI-based segmentation algorithm. Following segmentation, a custom ImageJ plugin (Labels to ROIs) was applied to extract the minimum Feret diameter (MinFeret) of individual fibers. The 5–7 measurements per mouse were subsequently averaged and plotted. To quantify CF ciliation, PDGFRα^+^ CFs co-labeled with the cilia marker ARL13B were counted across 3–5 fields of view using a 40x objective with a Z-resolution of 3–5 µm, and normalized to the total number of CFs within the corresponding area. Intramyocellular nuclei were subsequently identified using the Image Calculator function and quantified using the Analyze Particles function in ImageJ.

### Expression Analysis

Heart tissue was homogenized in TRIzol (ThermoFisher Scientific, cat. no. 15596026) using a bead beater (TissueLyser LT, Qiagen, cat. no. 69980) at 50 Hz for 5 minutes, with one 5 mm stainless steel bead (Qiagen, cat. no. 69989) and three 2.8 mm metal beads (Precellys, cat. no. KT03961-1-101.BK). Following the manufacturer’s protocol, chloroform was added to isolate the upper aqueous, RNA-containing phase. RNA was subsequently purified using the RNeasy Mini Kit (Qiagen, cat. no. 74106) according to the manufacturer’s instructions. A total of 500–800 ng of RNA was reverse-transcribed into cDNA using the qScript cDNA Synthesis Kit (Quantabio, cat. no. 84003). RT-qPCR was performed in technical quadruplicates on a QuantStudio 6 Flex Real-Time 384-well PCR System (Applied Biosystems, cat. no. 4485694) using PowerUp SYBR Green Master Mix (ThermoFisher Scientific, cat. no. A25742). Fold changes were calculated using the 2^-ΔΔCT^ method and expression levels were normalized to the housekeeping genes *Hprt, Sra1* and *Pde12*. Primer sequences are provided in Supplementary Table 1.

### RNA-sequencing

Single-cell RNA sequencing (scRNA-seq) data from two published databases (Hill, M. C., et al. 2022. PMID: 35732239; McLellan, M. A., et al. 2020. PMID: 32795101) were re-analyzed to examine the expression of Hedgehog (Hh) signaling, adipogenesis, fibrogenesis, and myogenesis-associated genes in the context of congenital heart disease and chronic cardiac stress. To further interrogate these pathways at the transcriptomic level, bulk RNA sequencing was performed on cardiac tissue isolated from control and CF^no cilia^ mice, with a focus on Hh signaling, adipogenesis, and fibrogenesis-related gene expression. For bulk RNA sequencing, RNA was isolated from formalin-fixed cardiac tissue sections using the RNeasy FFPE Mini Kit (Qiagen, cat. no. 74106) according to the manufacturer’s instructions. Libraries were prepared using a rRNA depletion-based protocol and sequencing was performed at the Interdisciplinary Center for Biotechnology Research (ICBR) at the University of Florida on an Illumina NovaSeq 6000 platform, generating 150 bp paired-end reads. Differential expression analysis was performed using DESeq2. Gene Ontology (GO) enrichment analysis was performed using ShinyGO. Genes with an adjusted p-value < 0.05 and an absolute log_2_ fold change > 1 were considered differentially expressed.

### Statistics

Injured hearts in which the injury encompassed less than 50% of the total area were excluded from the study. The experimenter was blinded to experimental conditions until all data were collected. All data were graphed using GraphPad Prism (version 10) and are presented as mean ± SEM. For comparisons between two groups with a single variable, an unpaired two-tailed Student’s *t*-test was used. For comparisons among more than two groups with a single variable, a one-way ANOVA followed by Dunnett’s multiple comparisons test was applied. For analyses involving two variables, a two-way ANOVA followed by Tukey’s multiple comparisons test was performed. A *p*-value of less than 0.05 was considered statistically significant, denoted as follows: \**p* < 0.05, \*\**p* < 0.01, \*\*\**p* < 0.001, and \*\*\*\**p* < 0.0001. Echocardiographic functional data were analyzed using multiple *t*-tests.

### Study Approval

All animal experiments were conducted in accordance with ethical guidelines and were approved by the Institutional Animal Care and Use Committee (IACUC) at the University of Florida and by the IACUC at the University of California, San Francisco.

## Data Availability

Transcriptomic data for *Dhh, Shh, Ihh, Gli1* and *Arl13b* were extracted from publicly available datasets. Single-nucleus RNA-seq data from Hill et al. (2022) are available in the NCBI Gene Expression Omnibus (GEO) under accession GSE203274. Cardiac non-myocyte data from McLellan et al. (2020) are available via ArrayExpress under accession E-MTAB-8810. Bulk RNA-seq data are available on NIH GEO XXX.

## Acknowledgements

We thank the members of the Kopinke laboratory for critical discussions. This work was supported by the US National Institutes of Health (NIH) grants R01AR079449 and R01HL171050 to D.K. and R01AR054396 and R01HD089918 to J.F.R. This research was enabled in part by support provided by the Chan Zuckerberg Biohub. The content is solely the responsibility of the authors and does not necessarily represent the official views of the NIH. This work is, in part, the result of NIH funding and is subject to the NIH Public Access Policy.

